# Modulation of *α*-Synuclein Aggregation Amid Diverse Environmental Perturbation

**DOI:** 10.1101/2023.10.19.563053

**Authors:** Abdul Wasim, Sneha Menon, Jagannath Mondal

**Affiliations:** Tata Institute of Fundamental Research, Hyderabad, India-500046

## Abstract

Intrinsically disordered protein *α*-Synuclein (*α*S) is implicated in Parkinson’s disease due to its aberrant aggregation propensity. In a bid to identify the traits of its aggregation, here we computationally simulate the multi-chain association process of *α*S in aqueous as well as under diverse environmental perturbations. In particular, the aggregation of *α*S in aqueous and varied environmental condition led to marked concentration differences within protein aggregates, resembling liquid-liquid phase separation (LLPS). Both saline and crowded settings enhanced the LLPS propensity. However, the surface tension of *α*S droplet responds differently to crowders (entropy-driven) and salt (enthalpy-driven). Conformational analysis reveals that the IDP chains would adopt extended conformations within aggregates and would maintain mutually perpendicular orientations to minimize inter-chain electrostatic repulsions. The droplet stability is found to stem from a diminished intra-chain interactions in the C-terminal regions of *α*S, fostering inter-chain residue-residue interactions. Intriguingly, a graph theory analysis identifies *small-world-like networks* within droplets across environmental conditions, suggesting the prevalence of a consensus interaction patterns among the chains. Together these findings suggest a delicate balance between molecular grammar and environment-dependent nuanced aggregation behaviour of *α*S.

## Introduction

In the human body, a significant presence of Intrinsically Disordered Proteins (IDPs) plays diverse and crucial roles.^1–3^ These proteins lack a well-defined 3D structure under native conditions, which imparts functional advantages, but also renders them susceptible to irreversible aggregation, especially when affected by mutations. Such aggregates can be pathogenic and are associated with various diseases, including neurodegenerative diseases, cancer, diabetes, and cardiovascular diseases. ^4^

Notably, Alzheimer’s disease is characterized by the aggregation of the amyloid-*β* peptide (A*β*), while Parkinson’s disease (PD) is linked to *α*-Synuclein (*α*S) aggregation. A growing body of evidence has established a connection between IDPs and the phenomenon known as *liquid-liquid phase separation* (LLPS). During LLPS, high and low concentrations of biomolecules coexist without the presence of membranes and exhibit properties similar to phase-separated liquid droplets of two immiscible liquids (Figure 1).^5–9^ This intriguing phenomenon has garnered significant attention as it underlies the formation of membrane-less subcellular compartments,^10–12^ which, when dysregulated, can lead to incurable pathogenic diseases.

**Figure 1.**
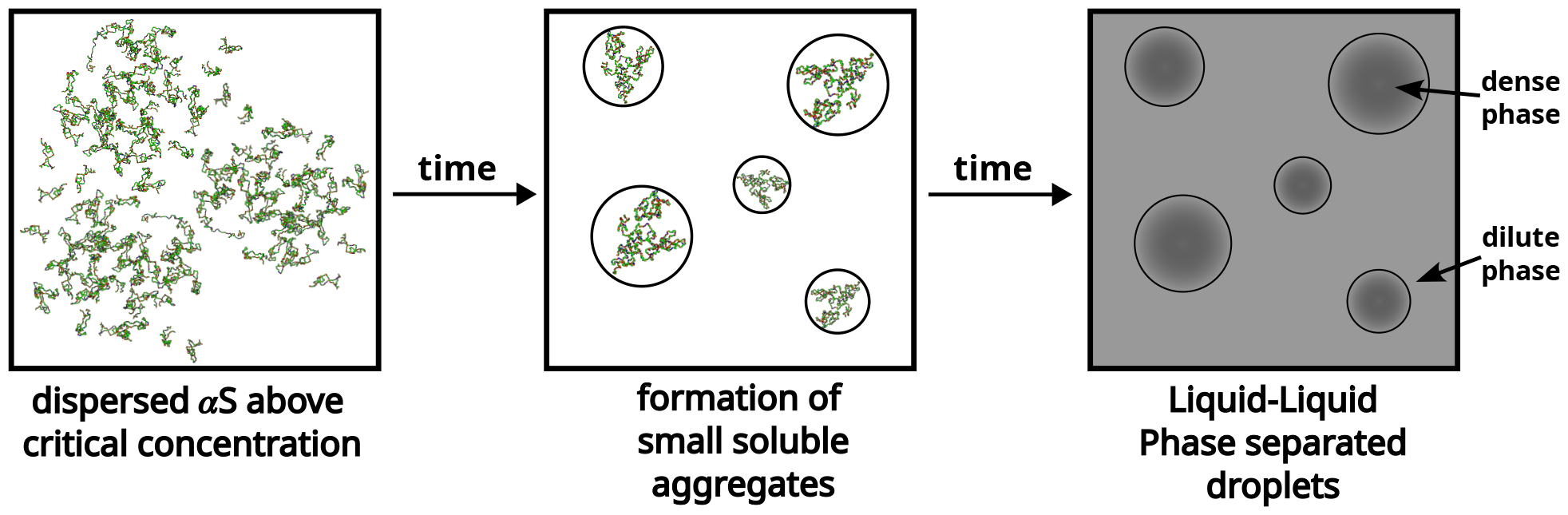
A schematic showcasing the process of liquid-liquid phase separation of *α*S.

Recent findings have highlighted the capability of *α*S to undergo LLPS under physiological conditions, specifically when the protein concentration surpasses a critical threshold.^6^ Moreover, it was observed that the aggregation propensity of *α*S is significantly influenced by various factors, including the presence of molecular crowders, the ionic strength of the protein environment, and pH.^13^ Nonetheless, characterizing the interactions and dynamics of these small aggregates poses experimental challenges, leading to limited available reports on the subject.^14–17^

This investigation aims to establish the molecular basis of self-aggregation of *α*S and underlying process of LLPS under diverse environmental perturbations. In particular, to understand the influence of environmental factors on the inter-protein interactions within a phase separated droplet, we target to computationally simulate the aggregation process of *α*S under different conditions, emphasizing the roles of crowders and salt. While recent progress in computational forcefields and hardware has enabled the simulation of individual Intrinsically Disordered Proteins (IDPs) especially *α*S, using All-Atom Molecular Dynamics (AAMD),^18–24^ these simulations can be extremely time-consuming and resource-intensive, making multi-chain AAMD simulations, even with cutting-edge software and hardware, impractical. Therefore, to simulate the the aggregation process, we resort to coarse-grained molecular dynamics (CGMD) simulations. Multiple coarse-grained forcefields have been developed with the sole purpose of fast and accurate simulations of IDPs and LLPS.^25–28^ However these are implicit water, residue-level coarse-grained models. Therefore, here we leverage a tailored Martini 3 Coarse-Grained Force Field (CGFF)^29^ for *α*S and use it to dissect the inter-protein interactions governing stable aggregate formation and LLPS. By leveraging the CGFF framework and building upon the groundwork laid by prior studies,^30–32^ we have optimized water-protein interactions for *α*S. Our multi-chain microsecond-long CGMD simulations have resulted in comprehensive ensembles of significant protein aggregates spanning various scenarios.

As one of the key observations, our simulation unequivocally reveal LLPS-like attributes in the aggregates and show how these get modulated in presence of crowders and salt. The investigation unearths the intricate interplay of mechanical and thermodynamic forces in *α*S aggregation, achieved through meticulous data analyses. We elucidate the pivotal intra and inter-protein interactions governing LLPS-like protein droplet formation, unveiling the protein’s primary sequence’s role in aggregation. As would be shown in this article, a graph-based depiction of the droplet’s architecture represents the proteins within droplets as constituting dense networks akin to *small-world networks*.

## Results

In this study, we utilized the recently developed Martini 3^29^ coarse-grained model to simulate collective interaction of a large number of *α*S chains in explicit presence of aqueous media at various concentrations commensurate with *in vitro* conditions including the presence of crowders and salt. As Martini 3 was not originally developed for intrinsically disordered proteins (IDPs), we carefully optimized the protein-water interactions against atomistic simulation of monomer and dimer of *α*S, as detailed in the *Methods* section, to ensure compatibility with *α*S (see *Methods*).

Initially, we examined the impact of concentration on the protein’s aggregation by simulating copies of chains, maintaining a polydispersity of protein conformations of *α*S. In particular, three different conformations of *α*S (referred here as *ms*1, *ms*2 and *ms*3) with *R*_*g*_s (radius of gyration) ranging between collapsed and extended states (1.84-5.72 nm) at different concentrations, with a composition, as estimated in a recent investigation,^23^ were employed. First the chains were simulated for extensive period in a set of three protein concentrations, close to previous experiments.

### Simulations capture enhanced aggregation beyond a threshold concentrations of *α*S

We performed simulations of *α*S at various concentrations, namely 300 *μ*M, 400 *μ*M, 500 *μ*M and 750 *μ*M. We begin by analysing the aggregation behavior of *α*S. As shown in Figure 2, we observe that most chains do not aggregate at 300 and 400 *μ*M as characterized by the prevalence of high number of free monomers. The respective snapshots of the simulation indicate the presence of greater extent of single chains. Also, the chains that are not free, form very small oligomers of the order of dimer to tetramer (Figure 2).

**Figure 2.**
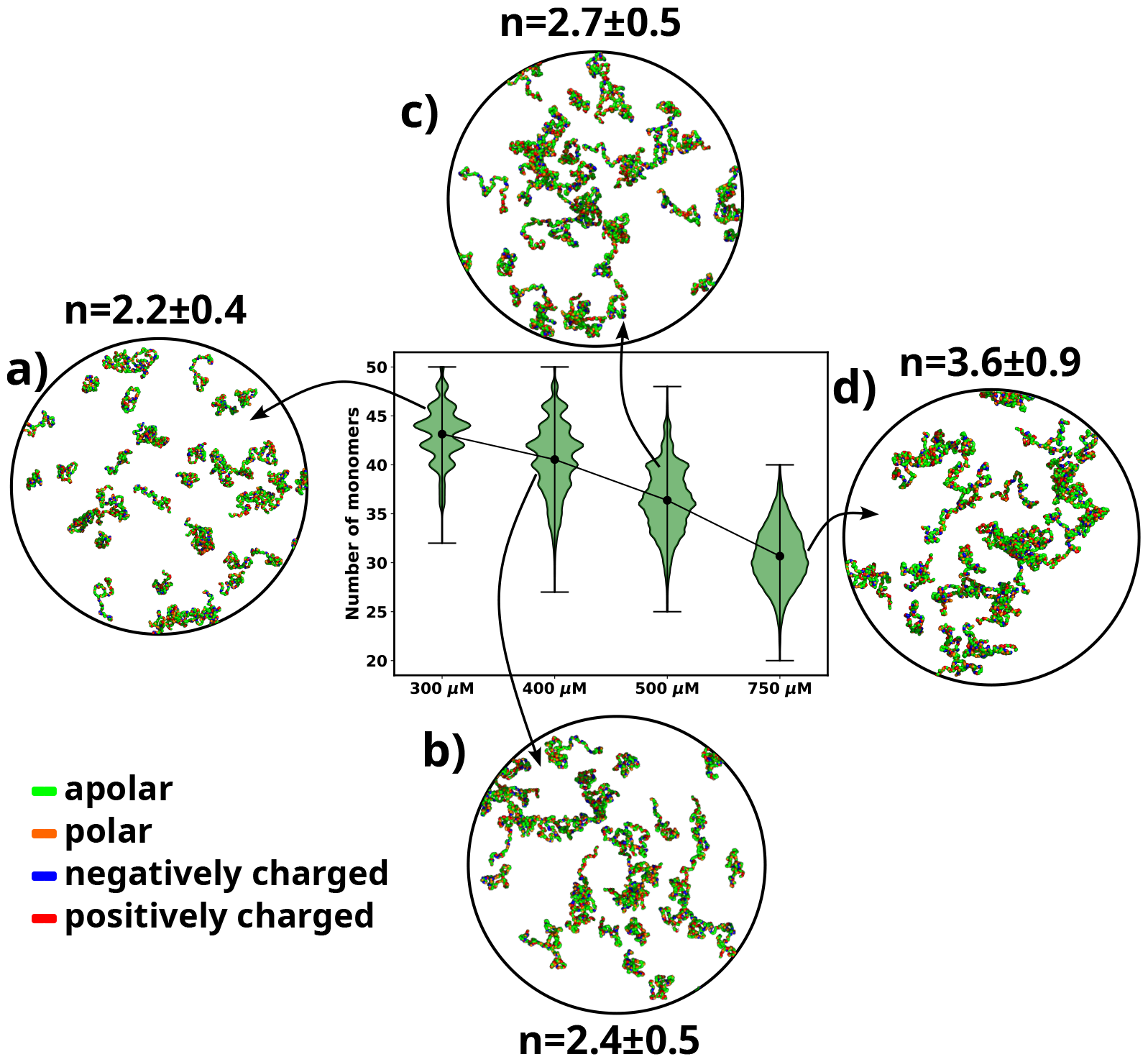
A violinplot showing the distribution of number of monomers present for different concentrations of *α*-syn. The blue dot at the middle of each distribution represents the mean number of monomers observed for each concentration. For each concentration we show representative snapshots of the system. For each concentration, we also report the statistics of the number of chains in the largest cluster (n). **a)** A snapshot from the simulation at 300 *μ*M *α*-syn. **b)** A snapshot from the simulation at 400 *μ*M *α*-syn. **c)** A snapshot from the simulation for 500 *μ*M *α*-syn. **d)** A snapshot from the simulation at 750 *μ*M *α*-syn. Each chain in the snapshots has been coloured differently.

However, upon increasing the concentration to 500 *μ*M, which has also been the critical concentration reported for *α*S to undergo LLPS,^6^ we observe a sharp drop in the average number of free monomers in the system (Figure 2). The corresponding representative snapshot of the system also depicts a few higher order aggregates, such as pentamers and hexamers, as well as most chains forming small oligomers. This can be understood from the value of the average number of chains present in the largest clusters, as reported in Figure 2.

The system, being at critical concentration, formation of large aggregates would require longer timescales than the simulation length. Therefore, in order to promote formation of large aggregates (heptamers or more) for finer characterization, we performed a simulation at a higher concentration of 750 *μ*M *α*S. As shown in Figure 2d, we observe further decrease in the total number of free monomeric chains in the solution. There is simultaneous appearance of a very few droplet like aggregates (hexamer or more) as can be seen from Figure 2 and the adjacent snapshot of the system (Figure 2). However, we note that ∼60 % of the protein chains are free and do not participate in aggregation and we think that as such in water, *α*S does not possess a strong and spontaneous self-aggregation tendency. In the following sections we characterize the aggregation tendency of *α*S in presence of certain environmental modulator that can shed more light on this hypothesis.

### Molecular crowders and salt accelerate *α*S aggregation

The cellular environment, accommodating numerous biological macromolecules, poses a highly crowded space for proteins to fold and function.^33–36^ In *in vitro* studies, inert polymers such as Dextran, Ficoll, and polyethylene glycol (PEG) are commonly employed as macro-molecular crowding agents. In the context of *α*S amyloid aggregation, previous experimental studies have revealed an increased rate of *in vitro* fibrillation in the presence of different crowding agents.^37–39^ Notably, a recent experimental study demonstrated the occurrence of phase separation (LLPS) of *α*S in the presence of PEG molecular crowder.^6^ Moreover, considering that *in-vivo* environments also contain various moieties like salts and highly charged ions, a recent *in vitro* study has shown that the ionic strength of the solvent directly influences the aggregation rates of *α*S,^13^ with higher ionic strength enhancing *α*S aggregation.

Given these observations, it becomes crucial to characterize the factors responsible for the enhanced aggregation of *α*S in the presence of crowders and salt. To address this, we perform two independent sets of simulations: one with *α*S present at 750 *μ*M in the presence of 10% (v/v) fullerene-based crowders (see *SI Methods*) and the other with the same concentration of *α*S but in the presence of 50 mM of NaCl. In this section we characterize the effects of addition of crowders or salt on the aggregation of *α*S.

As expected, the addition of crowders leads to an enhancement of *α*S aggregation due to their volume excluded effects, as depicted in Figure 3a. Notably, the number of monomers drastically decreases upon the inclusion of crowders. This observation is further supported by the snapshots of the system, which also confirm the reduction in monomer count. Similarly, we observe that the presence of salt also promotes *α*S aggregation, as illustrated in Figure 3a, where the number of monomers is lower when compared to the case with no salt.

**Figure 3.**
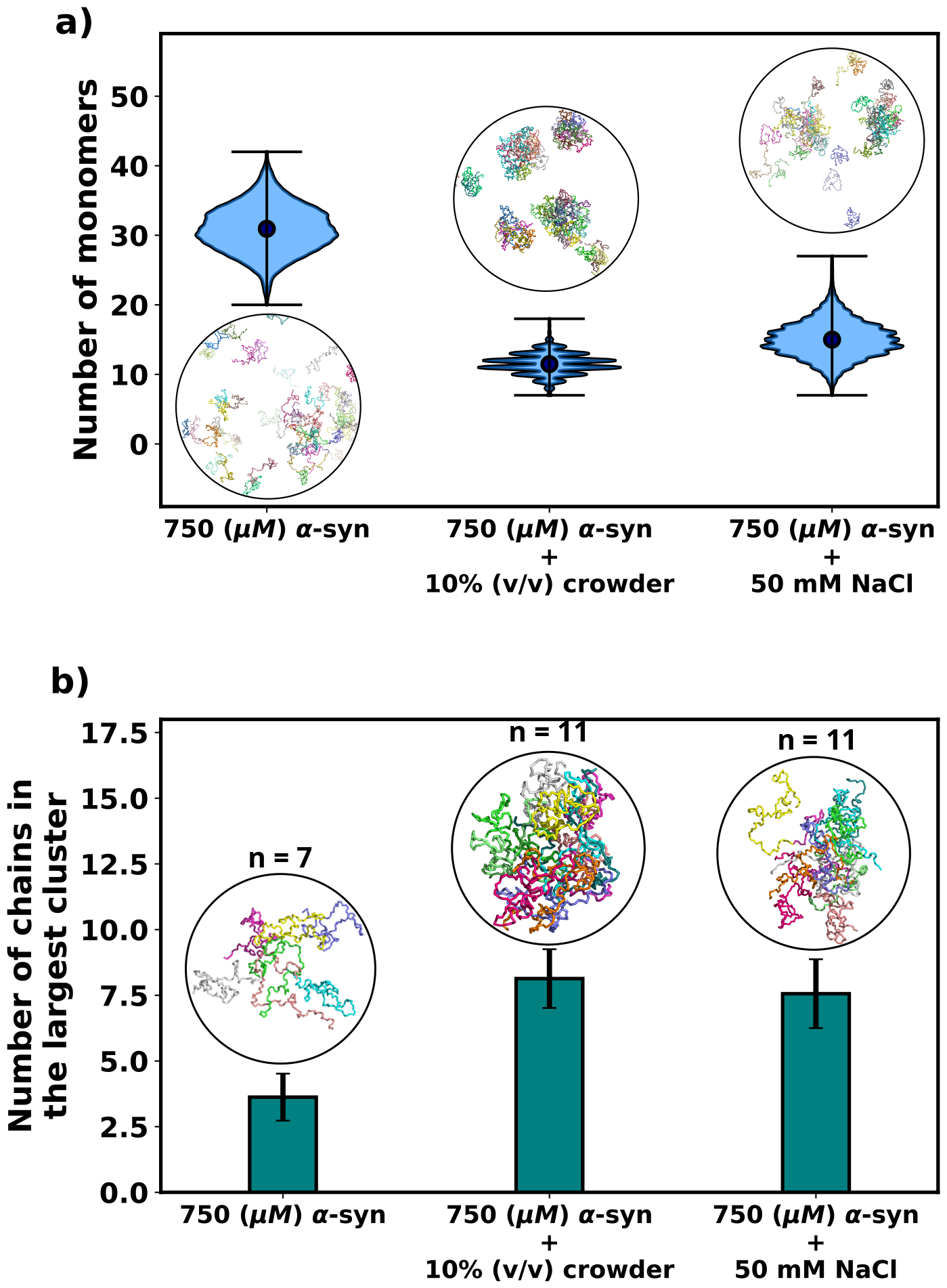
**a)** A violinplot showing the distribution of the number of monomers for *α*-syn at 750 *μ*M without and with crowder. The blue dots represent the means of each distribution. The snapshots represent the extent aggregation for a visual comparison. **b)** A bar plot showing the number of chain in the largest cluster formed by *α*-syn at 750 *μ*M without and with crowder. The snapshots show the largest cluster formed for each scenario.

Following this, we conducted an analysis of the number of chains present in the largest clusters that formed. Figure 3b clearly illustrates that the addition of crowder or salt leads to a notable increase in the average number of proteins forming a cluster. This crucial observation points to the fact that the inclusion of accelerators, such as crowder or salt, not only promotes aggregation but also plays a role in stabilizing the formed oligomers. Importantly, we observed that the effect of crowder on aggregation is slightly more pronounced compared to that of the salt. In the subsequent section, we delve into the reasons behind the enhanced aggregation induced by these accelerators, aiming to decipher the underlying mechanisms responsible for their influence on *α*S aggregation dynamics. As the aggregation is significant enough for performing quantitative analysis only when the concentration of *α*S is 750 *μ*M, we perform all analysis on scenarios at 750 *μ*M of *α*S.

### Crowders and salt differentially modulate surface tension for promoting LLPS-like *α*S droplets

The preceding sections underscore our simulation-based observation that, influenced by crowders and salt, *α*S aggregates into higher-order oligomers (hexamers and beyond) at a significantly accelerated propensity compared to the scenario without these influences. Here, we delve into the investigation of the energetic aspects underlying this aggregation phenomenon. An important contributor to the energetics is the surface tension, arising from the creation of interfaces between the dense and dilute phases of the protein upon droplet formation. This presence of interfaces is accompanied by surface tension and surface energy. The surface energy of a system is directly proportional to its surface area; systems with higher surface energy tend to minimize their surface area. Consequently, systems comprising multiple smaller droplets exhibit a larger surface area, and hence a higher surface energy. Conversely, systems characterized by fewer, larger droplets possess a comparatively reduced surface area and correspondingly lower surface energy. This insight leads us to conjecture that surface tension could play a pivotal role in driving liquid-liquid phase separation (LLPS) and the formation of larger *α*S droplets. To explore this hypothesis, we calculate the surface tension of the resultant droplets, as per Eqs-1, 2 and as described *SI Methods* and Ref.^30^

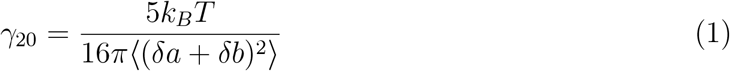

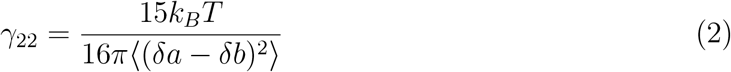

where *δa* = *a* − *R* and *δb* = *b* − *R* is the perturbation of the droplet shape from a perfect sphere with a radius *R* along any two pairs of principle axes of general ellipsoid estimating the shape of the droplet. The surface tension (*γ*) is thus estimated using *γ* ≈ *γ*_20_ ≈ *γ*_22_. Please see *SI Methods* and Ref.^30^ for more details.

Figure 4a provides a comparison of the surface tension (*γ*), for three different scenarios involving *α*S: i) *α*S in solution, ii) *α*S in the presence of crowders, and iii) *α*S in the presence of salt. Notably, in each case, the surface tension is considerably lower (0.0035-0.0075 mN/m) than the surface tension for FUS droplets in water (∼0.05 mN/m).^30^ As stated earlier, the magnitude of surface tension is an estimate of the aggregation tendency of any liquid-liquid mixture. Since we find that *γ*_*αS*_ is much lower than *γ*_*F US*_, we assert that the propensity with which *α*S aggregates should be much lower than that of FUS.

**Figure 4.**
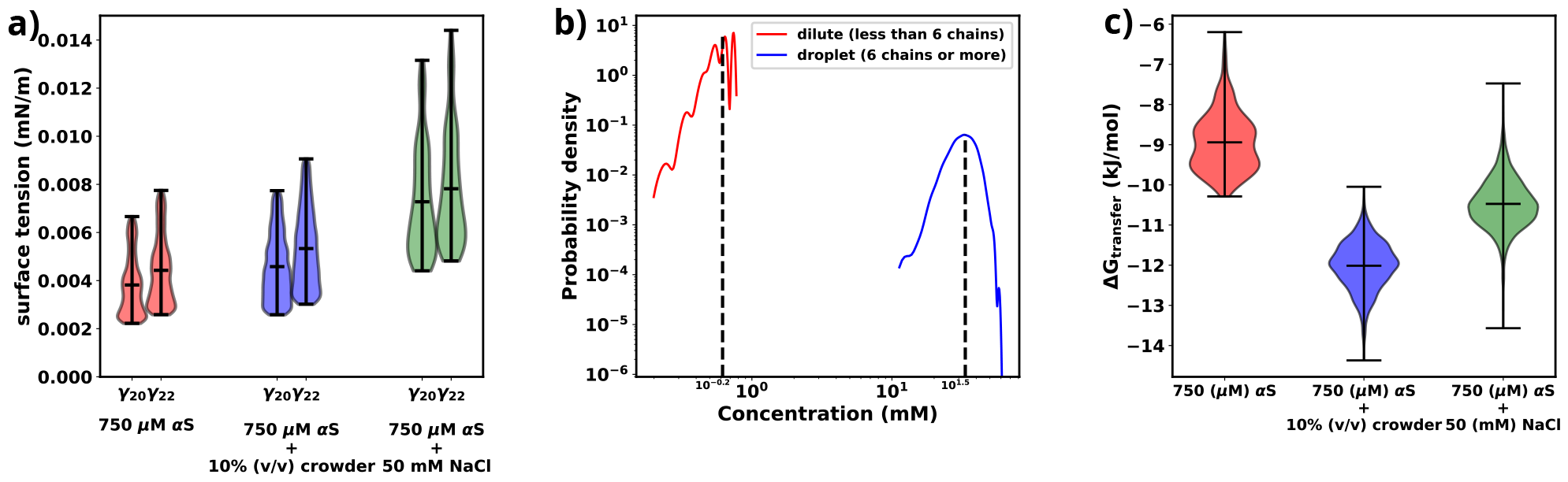
**a)** surface tensions of droplets, estimated from *γ*_20_ and *γ*_22_, for three cases has been shown. Both *γ*_20_ and *γ*_22_ provide almost similar estimates of the value of surface. **b)** Comparison of protein concentrations for the dilute(red) and the droplet (blue) phases. **c)** Excess free energy of transfer comparison for three cases.

Next, we conduct a comparison of the three different scenarios to understand the effects of crowders and salt on the aggregation of *α*S. From Figure 4a, it is evident that the surface tensions are very similar for cases (i) and (ii), while it has increased for case (iii). This implies that the addition of crowders does not significantly impact the surface tension of the aggregates, although it renders the protein more prone to aggregation. On the other hand, the addition of salt causes an increase in surface tension. Given the relationship between surface area and volume, where a higher surface-to-volume ratio signifies numerous smaller droplets, the surface energy is concurrently elevated. In the presence of salt, a tendency is observed for these smaller aggregates to coalesce, giving rise to larger aggregates, albeit in reduced numbers. This behavior is an endeavor to curtail the surface-to-volume ratio and thus mitigate the associated surface energy. Therefore, the larger the surface tension, the higher is tendency of the protein to form aggregates, as seen from the surface tension values of *α*S and FUS, as mentioned earlier.

To minimize the surface energy, fusion of aggregates, either via merging of two or more droplets into one is seen for liquid-like phase separated droplets in experiments.^6^ Although droplet fusion was not observed in our simulations due to the limited system size, it was shown that if a protein undergoes LLPS, a significant difference in protein concentration occurs between the droplet and the dilute phase.^40^ To verify whether the aggregates observed in our simulations exhibit characteristics of LLPS, we calculated the protein concentrations in the dilute and concentrated phases. For the droplet phase, the concentration of the protein was calculated using Eq-3.

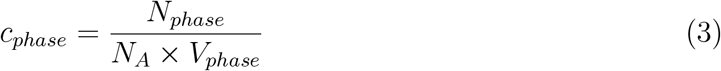

where *N*_*phase*_ is the number of protein chains in the phase (here dilute or concentrated), *N*_*A*_ is Avogadro’s number and *V*_*phase*_ is the volume occupied by the phase. For the dilute phase, we estimated the volume of the concentrated/dense phase (*V*_*dense*_) using Eq-4.^40^

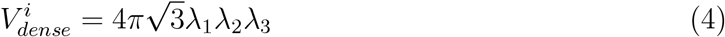

where 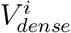 is the volume of the *i*-th droplet, *λ*_1_, *λ*_2_ and *λ*_3_ are the eigenvalues of the gyration tensor for the aggregate. The volume of the dilute phase is the remainder volume of the system given by Eq-5.

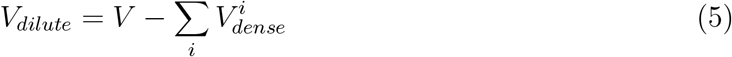

where V is the total volume of the system.

As shown in Figure 4b and Figure S1, there is an almost two orders of magnitude difference between the concentration of *α*S in the dilute and droplet phases for all scenarios. Such a pronounced difference is a hallmark of LLPS, leading us to assert that the aggregates formed in our simulations possess LLPS-like properties. Consequently, we use the term “droplet” interchangeably with “aggregates” for the remainder of our investigation.

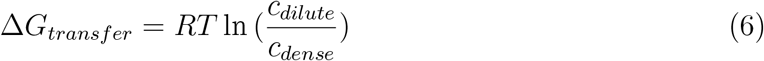

Finally, utilizing the calculated concentrations, we proceed to estimate the excess free energy of monomer transfer (Δ*G*_*transfer*_), from Eqn-6, between the dilute and droplet phases, where *c*_*dilute*_ is the concentration of *α*S in the dilute phase, *c*_*dense*_ is the concentration of *α*S in the dense/droplet phase, R is the universal gas constant and T is the temperature of the system (=310.15 K). As illustrated in Figure 4c, both crowder and salt scenarios demonstrate lower Δ*G*_*transfer*_ values compared to the case without their presence. However, the thermodynamic origins behind this pronounced aggregation differ for crowders and salt. Crowders enhance aggregation primarily through excluded volume interactions, which are of an entropic nature. On the other hand, salt enhances aggregation by increasing the droplet’s surface tension, thus contributing to the enthalpy of the system. As a result, apart from the already known fact that macromolecular crowding decreases Δ*G*_*transfer*_ via entropic means, we also infer that salt decreases Δ*G*_*transfer*_ via enthalpic means by increasing the surface tension of the formed droplets.

### Aggregation results in chain expansion and chain reorientation in *α*S

An indicative trait of molecules undergoing Liquid-Liquid Phase Separation (LLPS) is the adoption of extended conformations upon integration into a droplet structure. Given that the aggregates observed in our simulations exhibit a concentration disparity reminiscent of LLPS between the dilute and dense phases, we endeavored to validate the presence of a comparable chain extension phenomenon within our simulations.^40^ To address this, we quantified the radius of gyration (R_*g*_) for individual chains and classified them based on whether they were situated in the dilute or dense phase. The distribution of R_*g*_ values for each category is illustrated in Figure 5a and Figure S2. Remarkably, the distribution associated with the dense phase distinctly indicates that the protein assumes an extended conformation within this context. As elucidated earlier, this marked propensity for extended conformations aligns with a characteristic hallmark of LLPS as previously seen in experiments.^41^

**Figure 5.**
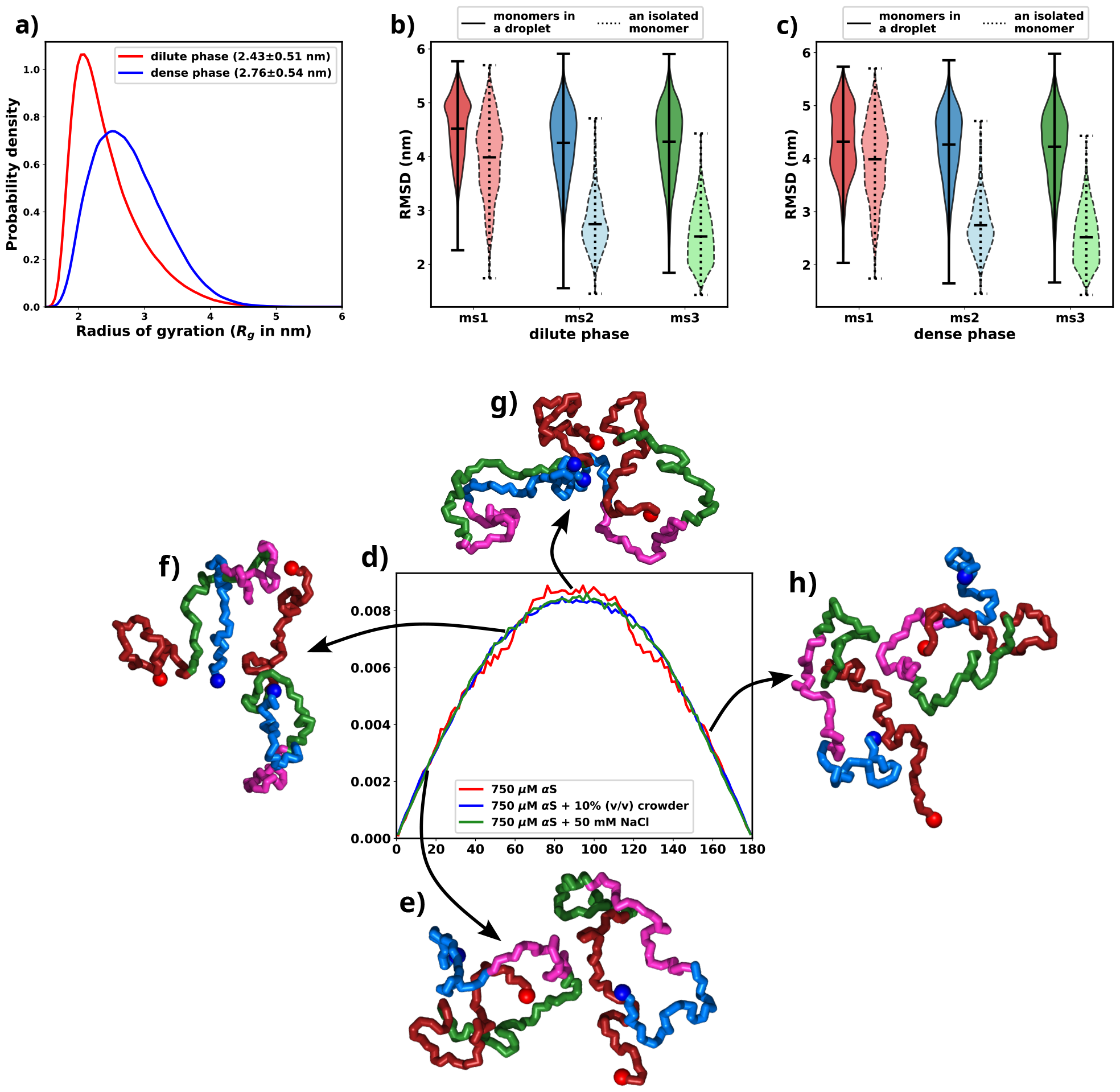
All the figures are for 750 *μ*M *α*S + 50 mM NaCl. **a)** Distribution of R_*g*_ for proteins present in the dense or the dilute phases. **b)** Comparison of RMSD for protein chains present in the dilute phase, with single chain RMSDs as the reference (dotted edges). **c)** Comparison of RMSD for protein chains present in the dense phase, with single chain RMSDs as the reference (dotted edges). **d)** Distribution of the angle of orientation of two chains inside the droplet for the three different scenarios. **e)** representative snapshot for angle between 0 degree and 20 degree. **f)** representative snapshot for angle between 50 degree and 70 degree. **g)** representative snapshot for angle between 80 degree and 120 degree. **h)** representative snapshot for angle between 150 degree and 180 degree.

Having observed the conformational alterations of *α*S during LLPS, our subsequent aim was to quantify the extent of these conformational changes in relation to their initial states (referred as ‘*ms*1’, ‘*ms*2’ or ‘*ms*3’ in decreasing order of *R*_*g*_ ^23^). To achieve this, we computed the Root Mean Square Deviation (RMSD) of each protein relative to its starting conformation. The resulting distributions were visually depicted using violin plots, featuring bold edges in Figures 5b and c. The protein ensemble was segregated into two categories: i) those from the dilute phase (Figure 5b), and ii) those from the dense phase (Figure 5c).

Surprisingly, regardless of their initial configurations, the observed RMSD values were notably high. To facilitate a comparative analysis, we also included distributions of RMSDs for single chains simulated in the presence of 50 mM of salt, depicted using violin plots with broken edges. Intriguingly, the conformational state labeled as *ms*1, exhibited the least RMSD, a characteristic attributed to its notably extended conformation. This phenomenon aligns with the preference of droplets for extended conformations, implying that ms1 required the least conformational perturbation and thus exhibited a lower RMSD.

For both ms2 and ms3, a conspicuous increase in RMSD values was observed across all proteins monomers, irrespective of their respective phases. This phenomenon can potentially be attributed to the pronounced conformational shift experienced by the protein during aggregation. Building on these observations, we put forward a hypothesis: LLPS engenders significant modifications in the native protein conformations, ultimately favouring the adoption of extended states.

As discussed in the previous paragraph that the *α*S monomers inside the droplets must undergo conformational expansion and we hypothesized that they adopt orientation so as to minimize the inter-chain electrostatic repulsions. To this end, we try to decipher the orientations of the chains via defining their axes of orientations and subsequently calculating the angles between the major axis of two monomers. We calculate the major axis of gyration, given by the eigenvector corresponding to the largest eigenvalue of the gyration tensor, for each monomer inside a droplet. We next find the nearest neighbour (minimum distance of approach *<* 8 Å) for each monomer, carefully taking care of over-counting.

The angle between two monomers is defined as the angle between the major axes of gyration between chain *i* and its nearest neighbour *j*. We plot the distributions of the angles for all scenarios and all droplets in Figure 5d. We observe that irrespective of the conditions, the distribution peaks at right angles. The representative snapshots (Figure 5e-5h) showcase their mode of orientation. Interestingly the distribution is the same for all the three scenarios, again stressing upon the fact the *α*S droplets share similar features in terms of interactions and orientations irrespective of their environments.

### Characterization of molecular interactions in aggregation-prone conditions

As established in preceding sections, both crowders and salt have been observed to augment the aggregation of *α*S while concurrently stabilizing the resultant aggregates. This phenomenon leads to the protein adopting extended conformations within a notably heterogeneous ensemble. Shifting our attention, we now delve into a residue-level investigation to unravel the specific interactions responsible for stabilizing these aggregates and, consequently, facilitating the aggregation process.

To compute the differential contact maps, our approach involved initial calculations of average intra-protein residue-wise contact maps, termed as intra-protein contact probability maps, for monomers present in both the dilute and dense phases (refer to Figures S3 and S4). Subsequently, we derived the difference by subtracting the contact probabilities of monomers within the dilute phase from those within the dense phase. As evident from Figure 6a, a discernible reduction in intra-chain Nter-Cter interactions is observed for monomers within the droplet phase, depicted by the presence of blue regions along the off-diagonals. Such a reduction in such interactions has also been observed via experiments^41^ and it is similarly noticeable in the two other cases, as evident in Figures 6b and 6c.

**Figure 6.**
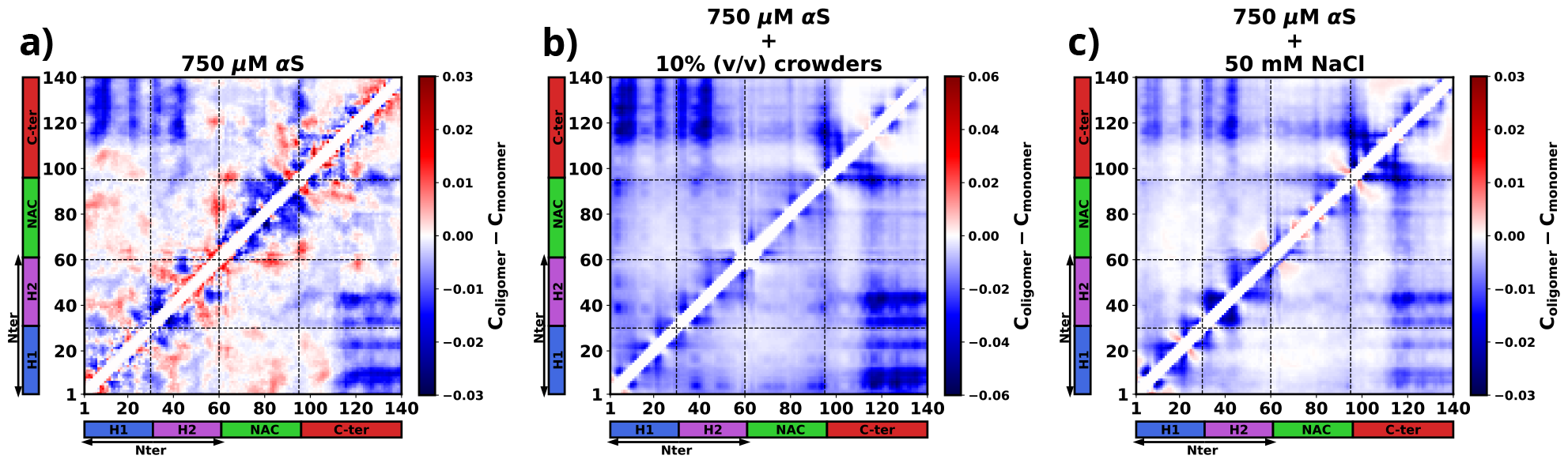
The figure presents the residue-wise, intra-protein difference contact maps where the average contact probability of monomers in the dilute phase were subtracted from the average contact probability of monomers in the dense/droplet phase for three cases: **a)** 750 *μ*M *α*S in water. **b)** 750 *μ*M *α*S in the presence of 10% (v/v) crowders. **c)** 750 *μ*M *α*S in presence of 50 mM NaCl.

Furthermore, a significant decline in intra-protein interactions, especially the NAC-NAC interactions, is predominantly observed at shorter ranges, indicated by deep-blue regions concentrated near the diagonals. Notably, these diminished intra-chain interactions (Figure 6 and S3) potentially facilitate the formation of inter-chain interactions (Figure S4). Thus we observed that increased inter-chain NAC-NAC regions (Figure S5) facilitate the formation of *α*S droplets which also have been previously been seen from FRET experiments on *α*S LLPS droplets.^6^ Building on these observations, we posit that these interactions play a pivotal role in stabilizing the aggregates that have formed.

Moreover, from the difference heatmaps in Figure S5, it can be observed that the residues 95-110 (VKKDQLGKNEEGAPQE) have reduced contact probabilities upon introduction of crowders/salt, whereas the rest of the contacts have slightly increased. These residues are highly charged and we think that upon introduction of crowders/salt, the proteins inside the droplet needed to be spatially oriented to facilitate the formation of largest aggregates. This re-orientation occurs to minimize the electrostatic repulsions among these residues belonging to different chains. These analyses provide hints that these residues are present in the protein so as to avoid the formation of aggregation prone conformations, which is why their interactions had to be minimized to form more stable and larger aggregates.

### Phase separated *α*S monomer form *small-world* networks

The investigations so far suggest that irrespective of the factors that cause the aggregation of *α*S, the interactions that drive the formation of droplet remain essentially the same. However the conformations of the monomers vary depending on their environment. In presence of crowders they adapt to form much more compact aggregates. Therefore here we characterize whether the environment influences the connectivity among different chains of the protein inside a droplet.

Figure 7a, 7b and 7c show molecular representations of the largest cluster formed by *α*S at 750 *μ*M in water, *α*S at 750 *μ*M in presence of 10% (v/v) crowders and *α*S at 750 *μ*M in presence of 50 mM NaCl respectively. From the molecular representations for aggregates, it can be seen that irrespective of the system, they form a dense network whose characterization is not possible directly. Therefore we represent each aggregate as a graph with multiple nodes (vertices) and connections (edges), as can be seen from Figure 7d, 7e and 7f. Each node (in blue) represents a monomer in the droplet. Two nodes have an edge (line connecting two nodes) if the minimum distance of approach of the monomers corresponding to the pair of nodes is at least 8 Å. We can see from the graph that not all chains are in contact with each other. They rather form a relay where a few monomers connect (interact) with most of the other protein chains. The rest of the chains have indirect connections via those. Since inter-chain connections/interactions have been denoted by edges and the chains themselves as nodes, such form of inter-chain interactions inside a droplet lead to only a few nodes having a lot of edges, for example node 1 in Figure 7f. The rest of them have only a few (3-5) edges. This is a signature of *small-world networks* ^42–45^ and we assert that *α*S inside the droplet(s) form small-world-like networks.

**Figure 7.**
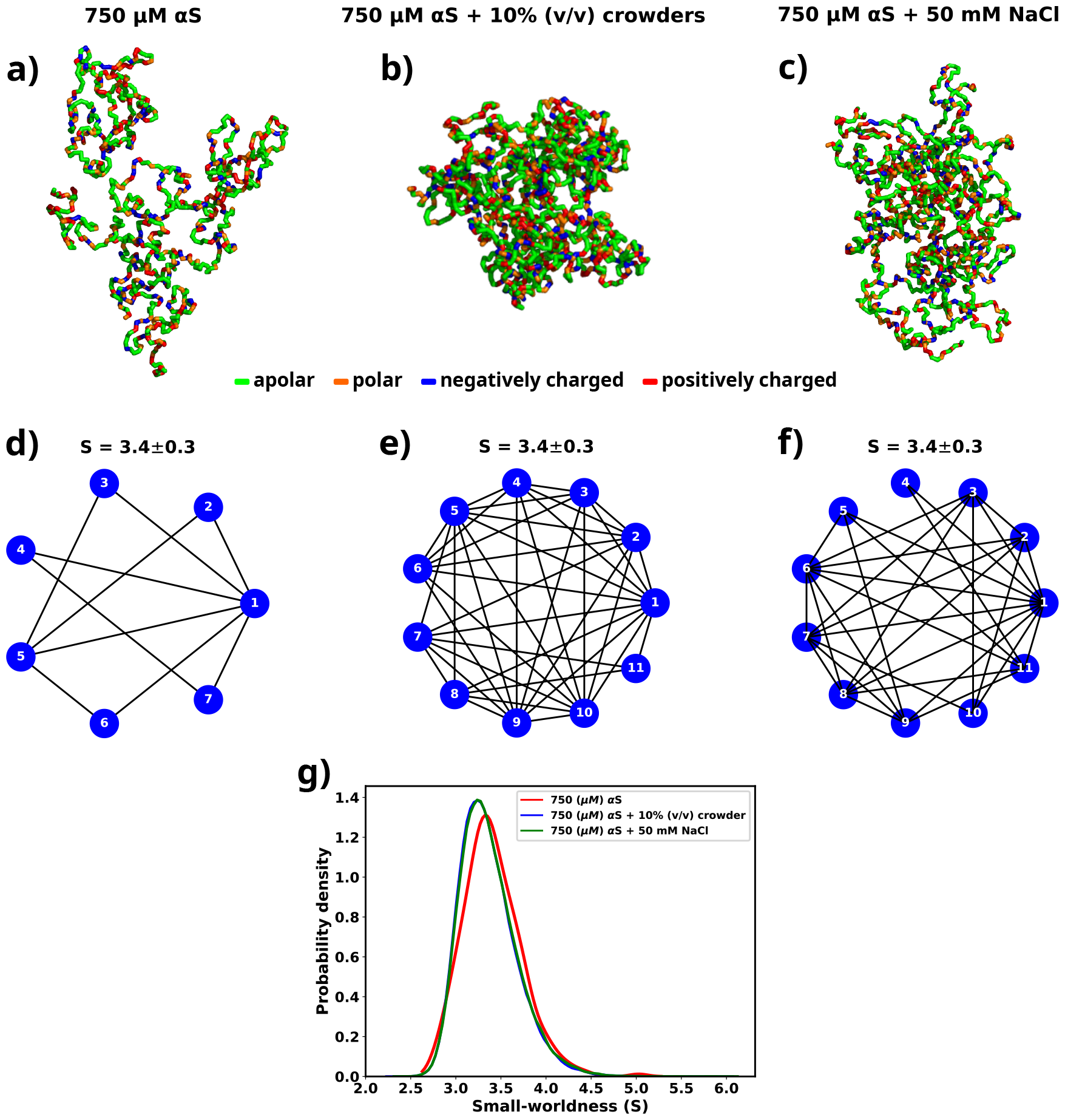
**a)** The largest cluster formed by *α*S at 750 *μ*M. **b)** The largest cluster formed by *α*S at 750 *μ*M in the presence of 10% (v/v) crowder. **c)** The largest cluster formed by *α*S at 750 *μ*M in the presence of 50 mM salt. Different residues have been color coded as per the figure legend. **d)** A graph showing the contacts among different chains constituting the largest cluster formed by *α*S at 750 *μ*M. **e)** A graph showing the contacts among different chains constituting the largest cluster formed by *α*S at 750 *μ*M in the presence of 10% (v/v) crowder. **f)** A graph showing the contacts among different chains constituting the largest cluster formed by *α*S at 750 *μ*M in the presence of 50 mM NaCl. The mean small-worldness (*S*) of all droplet has been reported above the graph. **g)** Distribution of small-worldness (*S*) for all scenarios.

A network can be classified as a small-world network by calculating the *Clustering coefficient* and the average shortest path length for the network and comparing those to an equivalent *Erdos-Renyi* network.^46^ The *clustering coefficient* (C) is a measure of the “connectedness” of a graph, indicating the extent to which nodes tend to cluster together. It quantifies the likelihood that two nodes with a common neighbor are also connected. On the other hand, the *average shortest path length* (L) is a metric that calculates the average number of steps required to traverse from one node to another within a network. It provides a measure of the efficiency of information or influence propagation across the graph. To estimate the *small-worldness* of a graph, we calculate a parameter (*S*) defined by Eq-7.

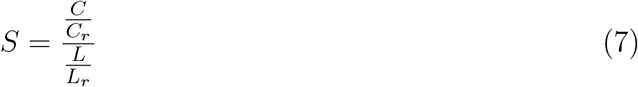

where *C* and *L* are the clustering coefficient and average shortest path length for the graph generated for a droplet while *C*_*r*_ and *L*_*r*_ clustering coefficient and average shortest path length for an equivalent Erdos–Renyi network. Small-world networks exhibit the characteristic property of having *C >> C*_*r*_, while *L* ≈ *L*_*r*_. In light of this, for every scenario (solely *α*S, *α*S in the presence of crowder, and *α*S in the presence of salt), we generate an ensemble of graphs that correspond to the droplets formed during the simulation.

For each graph, we calculate the small-worldness coefficient (*S*)^44^ and illustrate the distribution in Figure 7g. We observe a narrow distribution of *S* with a mean of 3.4 for all cases. In a previous report of RNA-LLPS, a value of *S* ≈ 4 was used to classify the droplets small-world networks.^40^ Therefore, *S* = 3.4 would suggest that the droplets formed during the simulations are small-world like. Moreover, we observe that the distribution of *S* is invariant with respect to the environment of the droplet.

Therefore we establish that the modes of interactions, orientations and even connectivities among *α*S monomers inside a droplet remain same even when their environments are extremely different. We think that this occurs since the residue-level interactions among different monomers inside the droplet are similar irrespective of the environment, as shown in a previous section. This puts forth a very interesting way of viewing *α*S LLPS. We think that if these residue-level interactions can be disturbed then the stability of the formed droplets might be affected in such a way that they might dissolve spontaneously.

## Discussion

We used simulations to investigate the molecular basis of *α*S monomeric aggregation into soluble oligomers resembling micro-LLPS. The WT protein demonstrated limited aggregation, suggesting a low inherent propensity for LLPS dictated by its primary sequence. IDPs, like *α*S, often share primary sequence characteristics associated with phase separation. Charged residues distributed with uncharged amino acids, resembling the “sticker and spacer” model, contribute to this molecular grammar. This observation aligns with a general trend in IDPs.^47–49^ To assess *α*S LLPS propensity from its primary sequence, we calculated Shannon entropy (*S*)^50^ (Eq 8 and Table S2), Kyte-Doolittle hydrophobicity^51^ (Table S3), Normalized, maximum of the sum of PLAAC Log-Likelihood Ratios (NLLR)^52^ (Table S4), and LLPS propensity scores obtained from catGranules web-server^53^ (Table S5).

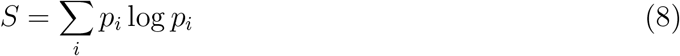

where *p*_*i*_ is the probability of occurance of a residue in a given sequence.

Comparative analysis with three datasets,^54^ namely LLPS+: a dataset of high propensity IDPs whose critical concentrations are 100 *μ*M or below, LLPS-: a dataset of low propensity IDPs whose critical concentrations are greater than 100 *μ*M, and PDB*: a dataset of folded proteins that do not undergo LLPS under normal conditions, revealed *α*S’s distinctive features (Table S6).

We note a significant difference in the Shannon entropy value of *α*S compared to proteins that do not undergo phase separation, as illustrated in Figure 8a. This deviation suggests a notable inclination of *α*S to undergo phase separation.^54^ Additionally, the hydrophobicity of *α*S (Figure 8b) is lower than that of the PDB* dataset, aligning more closely with the upper extremes of the LLPS-dataset. This indicates that while *α*S exhibits a tendency to undergo phase separation, the propensity should be low. Consistent with this, NLLR scores obtained from PLAAC and LLPS propensity scores (Figures 8c and d) reinforce this observation. These collective comparisons, coupled with simulations and experimental data on its critical concentration,^6^ conclusively establish that *α*S does not possess a high LLPS-forming propensity. Instead, this behavior is inherent to its primary structure. In hindsights, this analysis also justifies the requirements of environmental factors for enhancing the proclivity of *α*S for LLPS, as demonstrated in both our simulations and experimental findings.^6,13^

**Figure 8.**
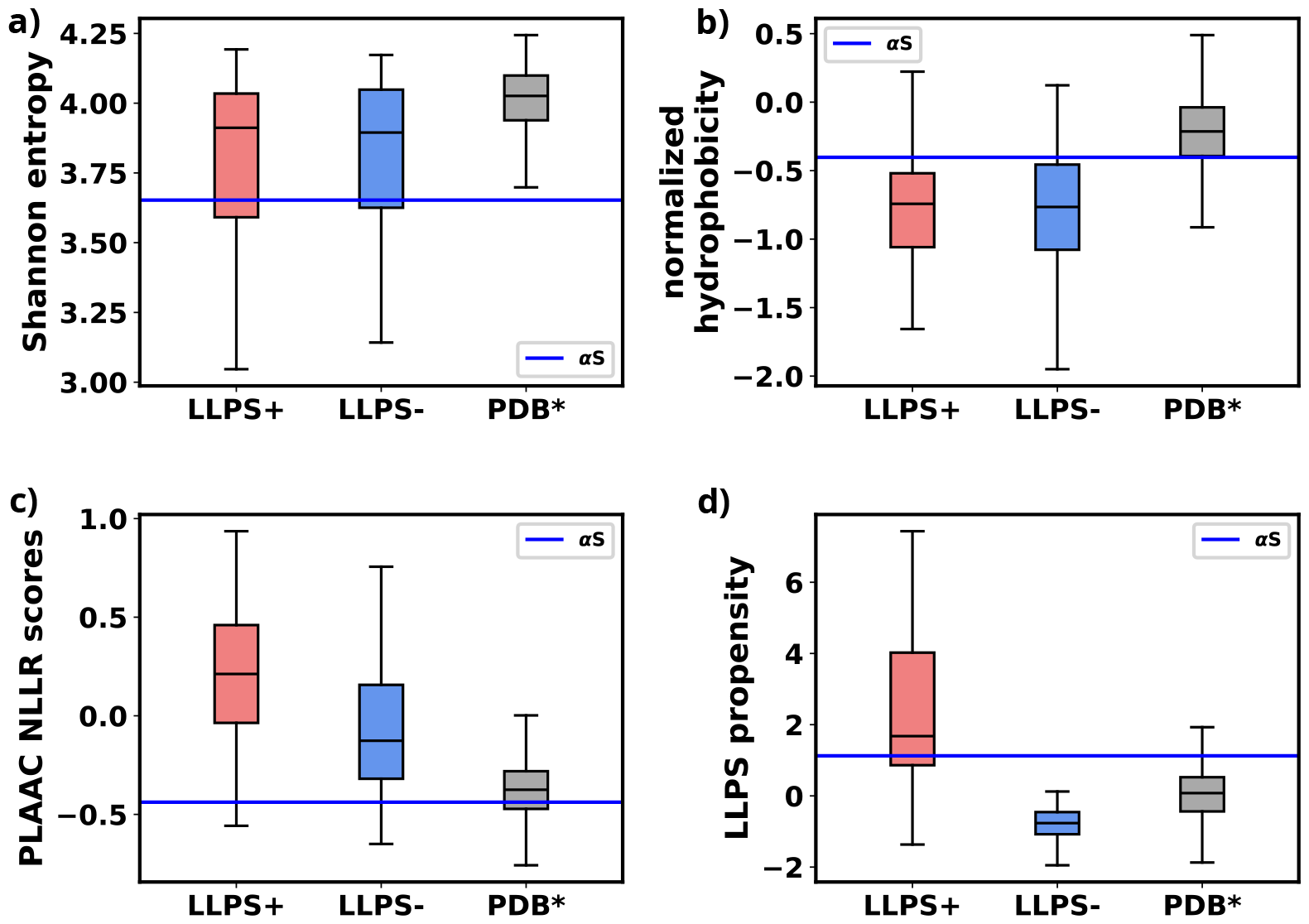
**a)** Comparison of Shannon Entropy of different datasets with *α*S. **b)** Comparison of Kyte-Doolittle hydrophobicity of different datasets with *α*S. **c)** Comparison of LLR scores, obtained from PLAAC, of different datasets with *α*S. **d)** Comparison of LLPS propensity scores, obtained from catGRANULE websever, of different datasets with *α*S. The values have been summarized in Table S6.

For characterizing *α*S’s aggregation phenomena, we calculated droplet surface tension under varied conditions. We observed that crowders minimally impacted surface tension, while salt increased it; however, both scenarios decreased the relative free energy of the system. Crowders achieved this via entropic means, whereas salt employed enthalpic means. Residue-residue interactions during droplet formation were consistent across environments, with crowder or salt enhancing these interactions. The aggregation pathway involved overall inter-chain interaction enhancement, specifically reducing intra-chain Nter-Cter and Nter-NAC interactions, leading to more extended protein conformations in droplets. The comparison with reported FRET observations^6^ aligns well with the findings from our simulations, indicating that within the droplets, intra-chain NAC-NAC interactions have been supplanted by inter-chain NAC-NAC interactions. Droplet proteins displayed consistent orientation and “small-worldness”, a measure of inter-chain connectivity, remained consistent across diverse conditions. Thus, *α*S aggregates appeared invariant regarding their initial environment in terms of interactions and contacts.

Our study’s precision was notably influenced by the careful selection of a simulation force-field. Despite the availability of modern force-fields optimized for multi-chain simulations of IDPs,^25,26,55,56^ we opted for Martini 3, an explicit water model, due to its emphasis on water’s role in aggregation and LLPS, as recently demonstrated in FUS LLPS.^57^ Although newer models operate at a faster pace, Martini 3’s inclusion of explicit water enhances result accuracy. Additionally, Martini 3 provides a detailed amino acid description and allowing for encoding of protein secondary structures, unlike some newer models that represent amino acids as single beads. Our meticulous choice of the simulation model, combined with a comprehensive analysis, contributes to the accuracy and novelty of this study.

Recent studies have explored the aggregation and LLPS of biopolymers and polyelectrolytes in the presence of membranes, opening a promising avenue for *α*S research.^58–60^ Given that under physiological conditions, *α*S assumes an oligomeric, membrane-bound form, investigating its interactions with membranes could hold therapeutic potential.^61^

Under physiological conditions, crowding effects emerge prominently. While crowders are commonly perceived to be inert, as has been considered in this investigation, the morphology, dimensions, and chemical interactions of crowding agents with *α*S in both dilute and dense phases may potentially exert considerable influence on its LLPS. Hence, a comprehensive understanding through systematic exploration is another avenue that warrants extensive investigation.

Although we exclusively focused on wild-type *α*S, familial mutations have been reported to exhibit a significantly higher propensity for aggregation.^6^ These mutations, involving minor alterations in the primary sequence, highlight the importance of understanding the molecular basis of this distinctive phenotype. Additionally, the observed stability of preformed *α*S droplets^62^ poses a challenge in treating Parkinson’s Disease (PD). Reversing aggregation/LLPS and understanding associated pathways and mechanisms are crucial. Our study identifies key residues crucial for stable droplet formation, consistent across various environmental conditions.

The significance of the solvent in *α*S aggregation remains underexplored. Recent studies^30,57^ underscore the pivotal role of water as a solvent in LLPS. It suggests that comprehending the solvent’s role, particularly water, is essential for attaining a deeper grasp of the thermodynamic and physical aspects of *α*S LLPS and aggregation. By delving into the solvent’s contribution, researchers can uncover additional factors influencing *α*S aggregation. Such insights hold the potential to advance our comprehension of protein aggregation phenomena, crucial for devising strategies to address diseases linked to protein misfolding and aggregation, notably Parkinson’s disease. Future investigations focusing on elucidating the interplay between *α*S, solvent (especially water), and other environmental elements could yield valuable insights into the mechanisms underlying LLPS and aggregation. Ultimately, this could aid in the development of therapeutic interventions or preventive measures for Parkinson’s and related diseases.

## Methods

### Selection of the metric for optimizing water-protein interactions

We have opted to utilize the radius of gyration (Rg) of *α*S as the primary metric for optimizing water-protein interactions in Martini 3 for *α*S. To calibrate the Martini 3 force field, we employed 73 *μ*s of all-atom data obtained from DE Shaw Research. From a polymer physics perspective, modifying water-protein interactions entails altering the solvent characteristics surrounding the biopolymer. We believe that Rg serves as an effective metric in this context. Additionally, we focus on matching the distribution of Rg values rather than solely targeting the mean value. This approach implies that, at a molecular level, the CGMD simulations conducted with optimized water-protein interactions enable the protein to explore conformations present in the all-atom ensemble.

Furthermore, we conducted cross-validation by comparing the fraction of bound states in all-atom and CGMD dimer simulations. This we claim that *R*_*g*_ is good metric to be used for tuning of water-protein interactions in Martini 3.

### Optimizing Martini 3 parameters for *α*S

Martini 3^29^ was trained using DES-Amber^63^ that is an atomistic forcefield tuned for single-domain and multi-domain proteins. Therefore, the default parameters of the coarse-grained model is not suited for simulations of disordered proteins and reported to underestimate the global dimensions of these systems in addition to overestimating protein-protein interactions. Previous attempts to simulate IDPs have modified the Martini force field by tuning the water-protein interactions, specifically, *σ* and *ϵ* of Lennard-Jones interactions to render them suitable for modeling a specific IDP or all IDPs.^30,31,64^ Here, we follow a similar protocol, however instead of tuning only the *ϵ* part of the water-protein Lennard-Jones interactions, we refine both the *σ* and *ϵ* parameters of the water-protein interactions (Eq-9).

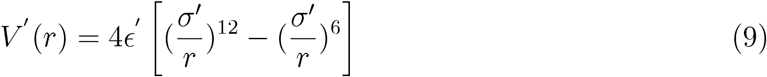

where *ϵ*^*′*^ = *λϵ, σ*^*′*^ = *λσ* and *λ* is the scaling parameter that needs to be optimized. Scaling *σ* tunes the relative radius of the hydration spheres of each residue of a protein while a change in *ϵ* changes the strength of the water-residue interactions (Figure 9a). Increasing the *ϵ* value of water-protein interactions results in a higher energy demand for removing water molecules (dehydration) as a chain transitions from the dilute to the dense phase. Conversely, a higher *σ* value implies that the hydration shell will be at a greater distance, facilitating dehydration if a chain moves into the dilute phase. Therefore, adjusting water-protein interactions based on the protein’s single-chain behavior may not significantly influence the protein’s phase behavior. Furthermore, fine-tuning both *ϵ* and *σ* parameters only requires a minimal change in the overall protein-water interaction (1%). As a result, this adjustment minimally alters the force field parameters.

**Figure 9.**
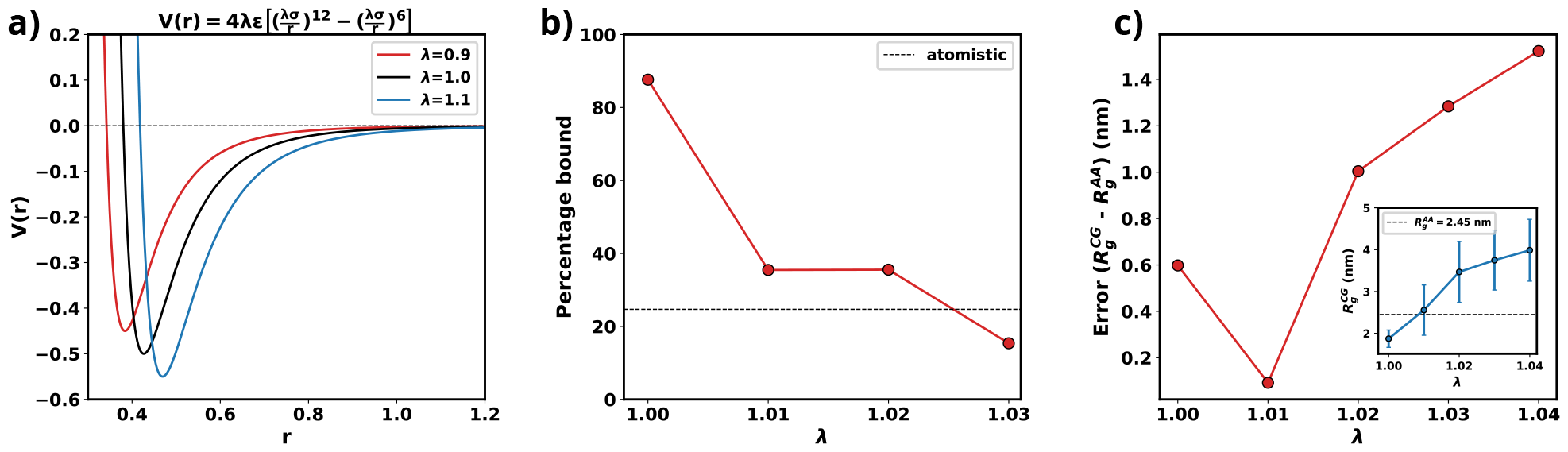
**a)** Plot of LJ potentials with respect to *λ*. **b)** The percentage bound values between two CG *α*S chains for different values of *λ*. The dashed black line represents the percentage bound values for two all-atom chains. **c)** Error between *R*_*g*_ calculated from CG and from all-atom simulations vs *λ*. The inset plot showcases the average values of *R*_*g*_ obtained from CG along with their respective standard deviations. The dashed line represents the average value from all-atom simulations.

As we are interested in exploiting multi-chain simulations to study the LLPS of *α*S using Martini 3, we use the percentage of time two all-atom monomers remain bound to each other as the benchmark. To obtain an optimum scaling parameter for the water-protein interactions in Martini 3, specific to *α*S, we perform CG simulations with two *α*S chains with different values of *λ*. We start with two chains, without any secondary structure enforced upon them, randomly placed in a 15.7 nm box making sure that they are apart by at least 0.8 nm which we use as cutoff to classify the chains to be bound. If the minimum distance between any two residues belonging to the different chains are closer than 0.8 nm we consider them to be bound. Using the cutoff defined, we calculate the percentage bound between the two *α*S monomers for different values of *λ* in the coarse-grained model. We also calculate the same from atomistic simulations reported in^23^ as the reference. From Figure 9b, we can see that for multiple values of *λ*, we observe a close agreement in percentage bound values between coarse-grained and atomistic simulations.

We conducted additional single-chain coarse-grained (CG) simulations of *α*S, varying the parameter *λ*, while refraining from imposing any secondary structure constraints. Subsequently, we compared the mean *R*_*g*_ values derived from these CG simulations with the 73 *μ*s all-atom (AA) trajectory, which replaced the previously published 30 *μ*s all-atom trajectory in^20^ and was provided by DE Shaw Research. Figure 9c illustrates that, for *λ* = 1.01, the average *R*_*g*_ in the CG simulations closely matches the *R*_*g*_ values obtained from the all-atom data. Consequently, we have chosen *λ* = 1.01 for the multi-chain simulations, as it minimizes errors for both single-chain *R*_*g*_ and the observed percentage of time bound in the two-protein chain simulations.

### Initial conformation generation for large-scale multi-chain simulations

A recent study used Markov State models to delineate the metastable states based on the extent of compaction (R_*g*_) and identified 3 macrostates and their relative populations.^23^ Subsequent to the investigation, we utilize three representative conformations, each corresponding to one of the macrostates. We designate these macrostates as 1 (ms1), 2 (ms2), and 3 (ms3) (refer to Figure S6). Therefore, in the multi-chain simulations, we maintain similar relative populations of these macrostates. (Figure S6). The reported percentages of macrostates (labeled as ms1, ms2 and ms3) are 0.06%, 85.9% and 14%, respectively. We added 50 *α*S monomers consisting of 1 chain of ms1, 45 chains of ms2 and 4 chains of ms3 in a cubic box with their respective secondary structures, determined via DSSP,^65–67^ enforced using Martini 3. The size of the box of side *a* is determined as per Eq-10.

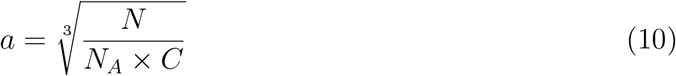

where *N* is the number of monomers, *N*_*A*_ is the Avogadro’s number and *C* is the required concentration of *α*S.

Here, we simulate multiple concentrations of the protein, namely, 300 *μM*, 400 *μM*, 500 *μM* and 750 *μM*. 300 and 400 *μM* are below the critical concentrations required to undergo LLPS.^6^ We then solvate the system in coarse-grained water. We setup the 50-chain system to simulate 3 conditions; (a) in pure water, (b) in 50 mM NaCl and (c) in presence of 10 % (v/v) crowders. To study effect of salt, we add the required number of Na^+^ and Cl^−^ ions to attain the desired concentration of 50 mM while also adding a few ions to render the system electrically neutral. In the system with crowders, we first add crowders after solvation by replacing a few solvent molecules with the required number of crowder molecules. We next resolvate the system along with the crowders. Finally we render the system electro-neutral by addition of the required number of Na^+^ or Cl^−^ ions. The details of the simulation setup are provided in Table 1.

**Table 1:** Details of the systems that were explored.

| summary | conc. of αS(μM) | box size (nm) | # water | # crowders | # $\text{Na}^+$ | # $\text{Cl}^-$ |
| --- | --- | --- | --- | --- | --- | --- |
| 300 μM αS in water | 300 | 65.66 | 2304122 | 0 | 450 | 0 |
| 400 μM αS in water | 400 | 59.69 | 1729213 | 0 | 450 | 0 |
| 500 μM αS in water | 500 | 55.52 | 1389721 | 0 | 450 | 0 |
| 750 μM αS in water | 750 | 48.42 | 920023 | 0 | 450 | 0 |
| 750 μM αS + 10%(v/v) crowder | 750 | 48.42 | 843011 | 20128 | 450 | 0 |
| 750 μM αS + 50 mM NaCl | 750 | 48.42 | 913357 | 0 | 3783 | 3333 |

\*For all simulations, a total of 50 monomeric protein chains have been used which comprise of $1 \times MS1$ , $45 \times MS2$ and $4 \times MS3$ .

### Simulation setup

Upon successful generation of the initial conformation, we first perform an energy minimization using steepest gradient descent using an energy tolerance of 10 kJ mol^−1^ nm^−1^. We next perform NVT simulations at 310.15 K using v-rescale thermostat for 5 ns using 0.01 ps as the time step. It is then followed by NPT simulation at 310.15 K and 1 bar using v-rescale thermostat and Berendsen barostat for 5 ns with a time step of 0.02 ps.

Next we perform CG MD simulations using velocity-verlet integrator with a time step of 0.02 ps using v-rescale thermostat at 310.15 K and berendsen barostat at 1 bar. Both Lennard-Jones and electrostatic interactions are cutoff at 1.1 nm. Coulombic interactions are calculated using Reaction-Field algorithm and relative dielectric constant of 15. We perform CG MD for at least 2.5 *μs* for the systems with 50 *α*S monomers. The details of the simulation run-times have been provided in Table S1. We use the last 1 *μs* for further analyses.

### Ascertaining the attainment of steady state in simulation

In this study, we utilized the final 1 *μ*s from each simulation for further analysis. To ascertain whether the simulations have achieved a steady state, we plotted the time profile of protein concentration in the dilute phase for all three cases.

Except for minor intermittent fluctuation involving only *α*S in neat water (Figures S7a and S7b), the remaining cases exhibit notably stable concentrations throughout various segments of the trajectory (Figures S7c-f). The relatively higher fluctuations observed in Figures S8a and b stem from the slow, spontaneous aggregation of *α*S alone, compounded by the inherently ambiguous nature of the dense phase. Consequently, the addition or removal of a few chains from the dense to the dilute phase results in significant fluctuations in protein concentration within the dilute phase. Conversely, in the other two scenarios (Figures S7c-f), aggregation is expedited by the presence of crowders/salt, leading to the formation of larger aggregates. Consequently, the addition or removal of one or two chains has negligible impact on concentration, thereby mitigating sudden large jumps. In summary, the conspicuous jumps depicted in Figures S8a and b arise from the gradual, fluctuating aggregation of pure *α*S and finite size effects. However, since these remain within the realm of fluctuations, we assert that the systems have indeed reached steady states. This assertion is bolstered by the subsequent figure, where the time profile of several pertinent system-wide macroscopic properties reveals no discernible change between 1.5-2.5 *μ*s (Figures S8).

### List of Software

We have used only open-source software for this study. All simulations have been performed using GROMACS-2021.^68,69^ Snapshots were generated using PyMOL 2.5.4.^70–72^ Analysis were performed using Python^73^ and MDAnalysis.^74,75^ Figures were prepared using Matplotlib,^76^ Jupyter^77^ and Inkscape.^78^

## Data Availability Statement

All data are present within the manuscript. The sections of the trajectories used for analysis (1.5 - 2.0 *μ*s) have been uploaded to zenodo (https://zenodo.org/records/10926368). Other relevant files related to the simulations have also been uploaded to the same repository. Additional data can be made available upon request.

## Acknowledgment

We acknowledge support of the Department of Atomic Energy, Government of India, under Project Identification No. RTI 4007. We also acknowledge Core Research grants provided by the Department of Science and Technology (DST) of India (CRG/2023/001426).

