## Supplementary material for "Modulation of *α*-Synuclein Aggregation Amid Diverse Environmental Perturbation": supplental text, supplemental table and figures

### Supplementary Information for Modulation of $\alpha$-Synuclein Aggregation Amid Diverse Environmental Perturbation

#### Supplementary Methods

##### Porting fullerene-based crowder to Martini 3

In this study, we model the crowders as fullerenes that have purely repulsive interactions with each other. Their interactions are modeled as consisting of only the repulsive part of their Lennard-Jones interactions instead of the full potential (Eq-1).

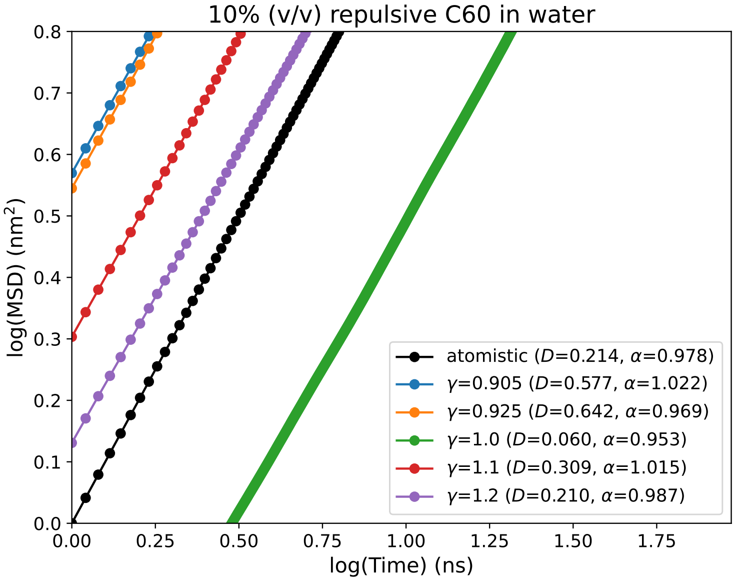


Figure 1: MSD vs time plots for different values of $\alpha$. The black line represents the MSD obtained from atomistic simulations with purely repulsive fullerene-fullerene interactions.

$$V^{F-F}(r) = \frac{4\gamma\epsilon\sigma^{12}}{r} \tag{1}$$

where $V^{F-F}(r)$ is the interaction among different fullerenes and $\gamma = 1.0$ for the default parameters reported for Martini 2.

The parameters previously reported for fullerene is for Martini 2 coarse-grained forcefield. Therefore, we port the parameters first to Martini 3 by addition of new interactions in Martini 3 force field (CNP beads). We test the validity of the ported parameters of fullerene by calculating and comparing their mean squared displacements (MSD) with those obtained from atomistic simulations (see Figure 1). For this, we performed atomistic simulation of 10 % (v/v) fullerene in water in a cubic box of ∼5 nm as it is the concentration used with $\alpha$S monomers as reported previously.[1] In a similar setup, we also run CG simulations of 10%(v/v) of fullerenes in water, where the volume of each fullerene based-crowder has been set to 0.55 nm$^3$.[2] As shown in Figure 1, the default ported parameters of fullerene do not reproduce the MSD obtained in atomistic simulations. This indicates that the fullerene parameters need to be tuned to obtain a good agreement in this dynamical property (MSD).

To achieve this, similar to the previous approach taken for modeling of $\alpha$S in Martini 3, we tune the water-CNP interactions in Martini 3 (Eq-1). We iteratively vary $\gamma$ to match the MSD from CG simulations to the reference atomistic one. We observe that at $\gamma = 1.2$, we obtain the closest match between Martini 3 CG and atomistic simulations (Figure 1).

#### Calculation of concentration of phases

We quantify the concentration of the protein in the solution (dilute phase) and in the aggregate (high-density phase) similar to the approach taken by Nguyen et al.[3] We first calculate the volume of aggregate by Eqn-2 as follows:

$$V_{aggr} = \frac{4\pi}{3}\sqrt[3]{\lambda_1\lambda_2\lambda_3} \tag{2}$$

where $\lambda_1$, $\lambda_2$ and $\lambda_3$ are the eigenvalues from the gyration tensor of the aggregate. The concentration of the proteins in the aggregate is then calculated by Eqn-3.

$$C_{aggr} = \frac{N}{N_A.V_{aggr}} \tag{3}$$

where $N$ is the number of chains in the aggregate, $N_A$ is the Avogadro's number and $V_{aggr}$ is the volume of the aggregate obtained using Eqn-2

The concentration of the dilute phase is then calculated by Eqn-4.

$$C_{dilute} = \frac{N_{dilute}}{N_A.V_{dilute}} \tag{4}$$

where $N_{dilute}$ is the number of chains present in trimers or lower aggregates, $N_A$ is the Avogadro's number and $V_{dilute} = V_{system} - \sum_i V^i_{aggr}$. $\sum_i V^i_{aggr}$ is the total volume occupied by larger aggregates (6 or more).

#### Calculation of small-worldness (S)

It is defined as per Eqn-5.[4]

$$S = \frac{\gamma_G}{\lambda_G} \tag{5}$$

where $\gamma_G = \frac{C_G^{\Delta}}{C_{rand}^{\Delta}}$ and $\lambda_G = \frac{L_G}{L_{rand}}$. $C_G$ is the mean clustering coefficient for a graph $G$ and $L_G$ is the mean shortest path length for $G$. $C_{rand}^{\Delta}$ and $L_{rand}$ are the mean clustering coefficient and mean shortest path length for an ensemble of Erdos-Renyi random network of the same size.

#### Calculation of surface tension

We follow the procedure reported by Benayad et al.[5] with only one minor difference. In[5] exact masses for the Martini beads were not taken into account, rather all beads were assumed to have the same mass. In this study we use the actual masses of the beads for all calculations.

As per,[5] we first calculate the droplet shape using Eqs-6 and 7.

$$C_{\alpha,\beta} = \frac{m_i(r_i^{\alpha} - r_{CMS}^{\alpha})(r_i^{\beta} - r_{CMS}^{\beta})}{\sum_i m_i} \tag{6}$$

where $C$ is the mass weighted covariance matrix, $\alpha$ and $\beta$ are directions $x, y$ or $z$; $i$ is the index for atoms/beads of protein monomers within a droplet, $r_{CMS}^{\alpha/\beta}$ is the centre of mass of the droplet in $x, y$ or $z$ direction; $m_i$ is the mass of the atom/bead.

The eigenvalues $\lambda_1, d_2$, and $\lambda_3$ of $C$ are given by: $\lambda_1 = vh^2, \lambda_2 = vb^2, \lambda_3 = vc^2$. Since $R^3 = abc$, where R is the average droplet radius, we obtain Eqs-7

$$a = \frac{R\lambda_1^{1/3}}{(\lambda_2\lambda_3)^{1/6}}$$

$$b = \frac{R\lambda_2^{1/3}}{(\lambda_1\lambda_9)^{1/6}}$$

$$c = \frac{R\lambda_3^{1/3}}{(\lambda_1\lambda_2)^{1/6}} \tag{7}$$

We next define $\delta a = a - R$, $\delta b = b - R$ and $\delta c = c - R$. Using these, we obtain Eq-8

$$\left\langle (\delta a \pm \delta b)^2 \right\rangle = \frac{1}{3} \sum_{i=1}^{2} \sum_{j=i+1}^{3} \left\langle (\delta a_i \pm \delta a_j)^2 \right\rangle \tag{8}$$

Therefore, the surface tension ($\gamma$) is them estimated using $\gamma \approx \gamma_{20} \approx \gamma_{22}$ where

$$\gamma_{20} = \frac{5k_BT}{16\pi\langle(\delta a + \delta b)^2\rangle}$$

$$\gamma_{22} = \frac{15k_BT}{16\pi\langle(\delta a - \delta b)^2\rangle} \tag{9}$$

#### Supplementary Tables

Table S1: Run-times of different simulations

| system | no. of replicas | runtime(s) |
|---|---|---|
| 300 $\mu$M $\alpha$S | 1 | 2.5 $\mu$s |
| 400 $\mu$M $\alpha$S | 1 | 4.3 $\mu$s |
| 500 $\mu$M $\alpha$S | 1 | 4.1 $\mu$s |
| 750 $\mu$M $\alpha$S | 4 | 2.6, 3.1, 3.0, 3.5 $\mu$s |
| 750 $\mu$M $\alpha$S + 10% (v/v) crowders | 4 | 2.8, 2.5, 2.6, 2.6 $\mu$s |
| 750 $\mu$M $\alpha$S + 50 mM NaCl | 4 | 2.6, 2.4, 2.6, 2.3 $\mu$s |

Table S2: Shannon entropy[6] for various datasets and $\alpha$S

| Dataset | min | mean | max |
|---|---|---|---|
| LLPS+ | 2.08 | 3.76 | 4.19 |
| LLPS- | 2.03 | 3.75 | 4.17 |
| PDB* | 3.08 | 4.00 | 4.24 |
| $\alpha$S | —— | 3.65 | —— |

Table S3: normalized Kyte Doolittle hydrophobicity[7] scores for various datasets and $\alpha$S

| Dataset | min | mean | max |
|---|---|---|---|
| LLPS+ | -2.19 | -0.75 | 0.95 |
| LLPS- | -2.16 | -0.78 | 1.05 |
| PDB* | -1.50 | -0.21 | 1.59 |
| $\alpha$S | —— | -0.41 | —— |

Table S4: PLAAC NLLR[8] scores for various datasets and $\alpha$S

| Dataset | min | mean | max |
|---|---|---|---|
| LLPS + | -0.558 | 0.206 | 0.936 |
| LLPS- | -0.650 | -0.035 | 0.936 |
| PDB* | -0.966 | -0.384 | 0.052 |
| $\alpha$S | —— | -0.438 | —— |

Table S5: catGRANULE[9] scores for various datasets and $\alpha$S

| Dataset | min | mean | max |
|---|---|---|---|
| LLPS+ | -1.37 | 2.37 | 9.83 |
| LLPS- | -2.03 | 1.43 | 8.26 |
| PDB* | -4.06 | 0.01 | 2.41 |
| $\alpha$S | —— | 1.13 | —— |

Table S6: Comparison of primary sequence derived features for various datasets and $\alpha$S

| Dataset | S | normalized hydrophobicity | NLLR | catGranule |
|---|---|---|---|---|
| LLPS+ | 3.76 | -0.75 | 0.206 | 2.37 |
| LLPS- | 3.75 | -0.78 | -0.035 | 1.43 |
| PDB* | 4.00 | -0.21 | -0.384 | 0.01 |
| $\alpha$S | 3.65 | -0.41 | -0.438 | 1.13 |

### Supplementary Figures

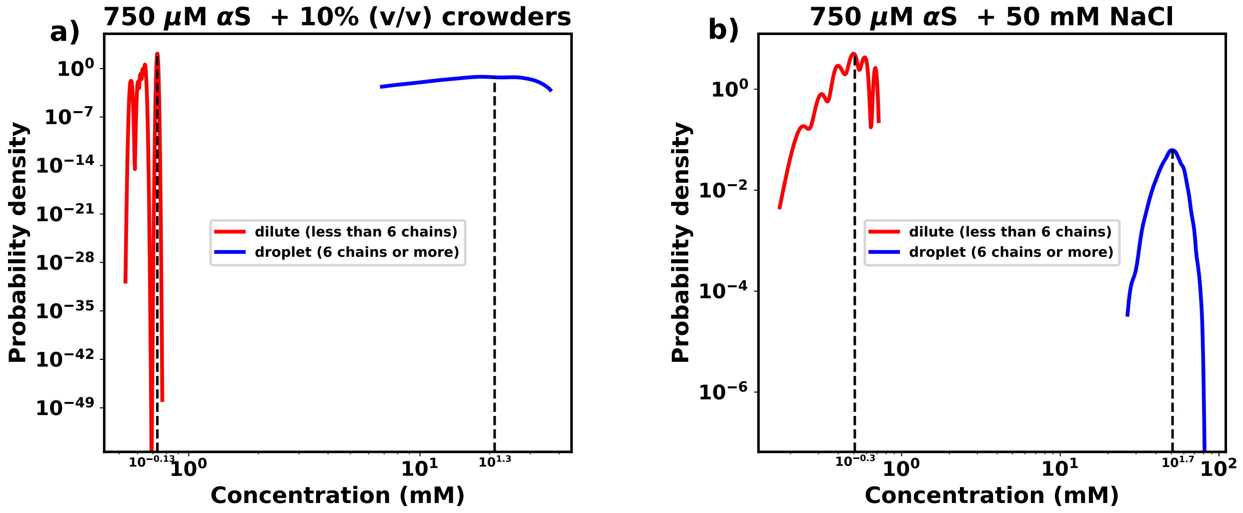


Figure S1: **a)** comparison of protein concentrations in the dense and the dilute phase for 750 $\mu$M $\alpha$S in the presence of 10% (v/v) crowders. **b)** comparison of protein concentrations in the dense and the dilute phase for 750 $\mu$M $\alpha$S in the presence of 50 mM NaCl.

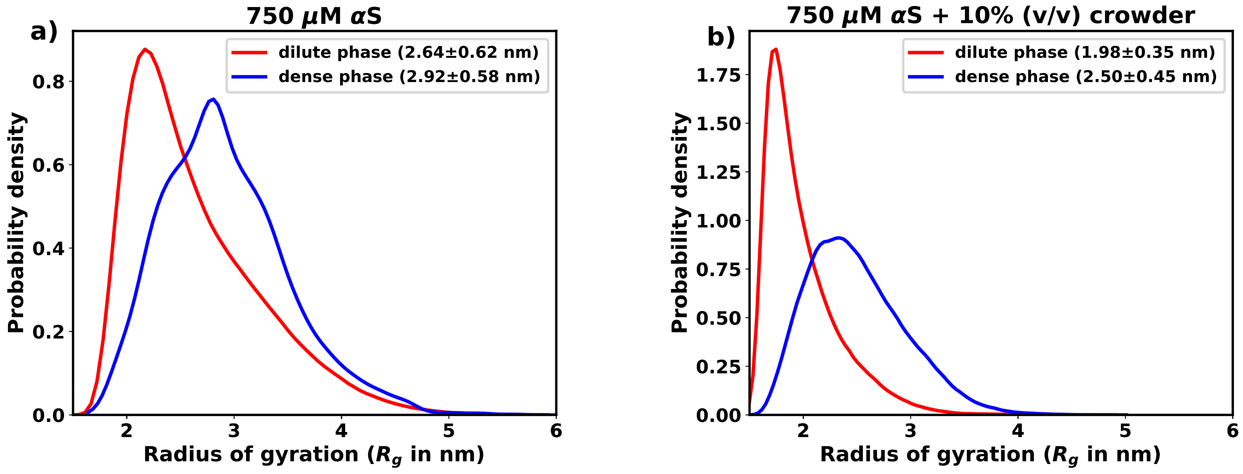


Figure S2: **a)** comparison of $R_g$ in the dense and the dilute phase for 750 $\mu$M $\alpha$S. **b)** comparison of $R_g$ in the dense and the dilute phase for 750 $\mu$M $\alpha$S in the presence of 10% (v/v) crowders.

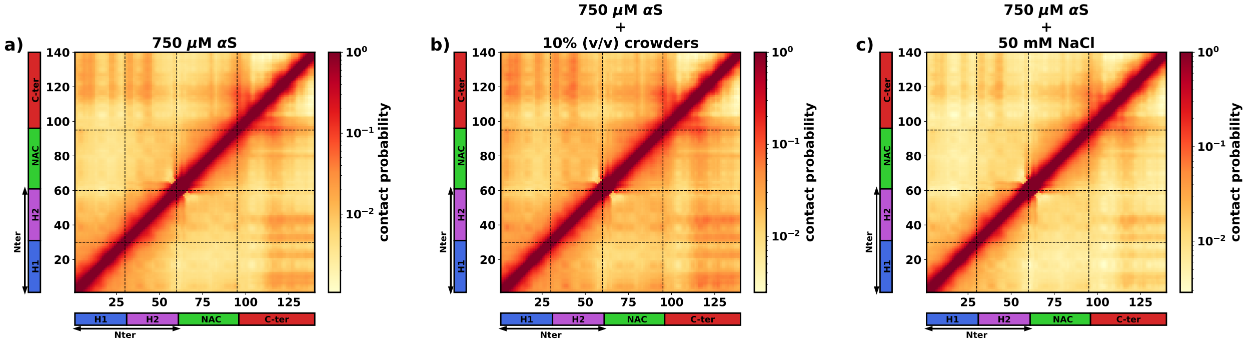


Figure S3: The intra-protein contact probability heatmap for proteins in the dilute phase for three scenarios: **a)** 750 $\mu$M $\alpha$S. **b)** 750 $\mu$M $\alpha$S + 10% (v/v) crowders. **c)** 750 $\mu$M $\alpha$S + 50 mM NaCl.

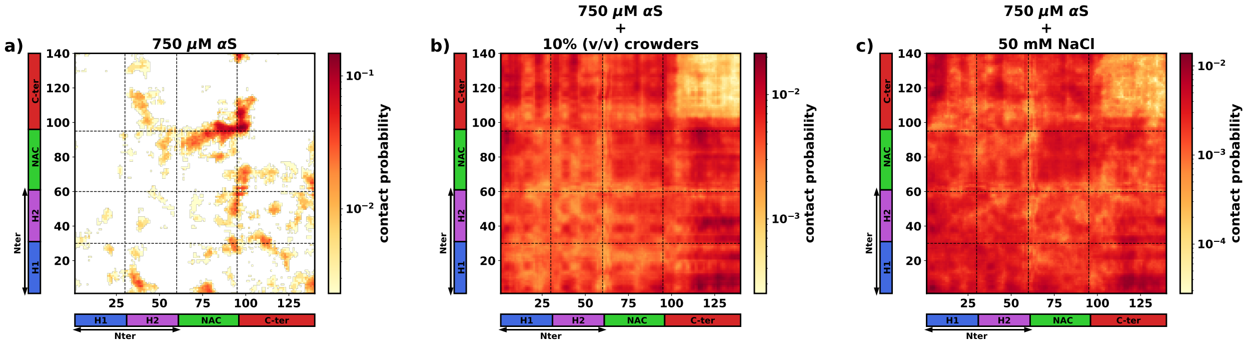


Figure S4: The inter-protein contact probability heatmap for proteins in the dense phase for three scenarios: **a)** 750 $\mu$M $\alpha$S. **b)** 750 $\mu$M $\alpha$S + 10% (v/v) crowders. **c)** 750 $\mu$M $\alpha$S + 50 mM NaCl.

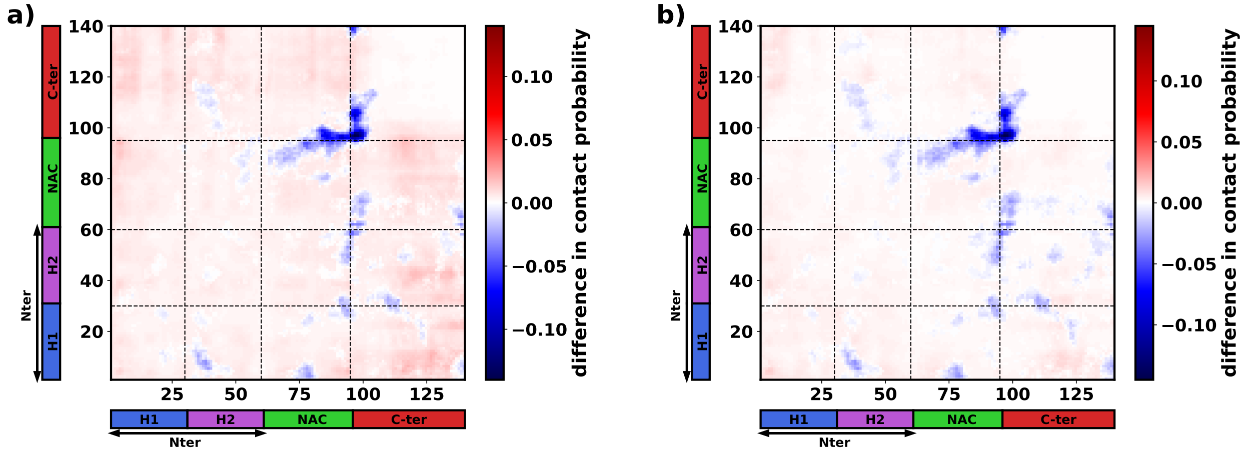


Figure S5: The difference in inter-protein contact probabilities heatmap for proteins in the dense phase. **a)** 750 $\mu$M $\alpha$S + 10% (v/v) crowders - 750 $\mu$M $\alpha$S. **b)** 750 $\mu$M $\alpha$S + 50 mM NaCl - 750 $\mu$M $\alpha$S.

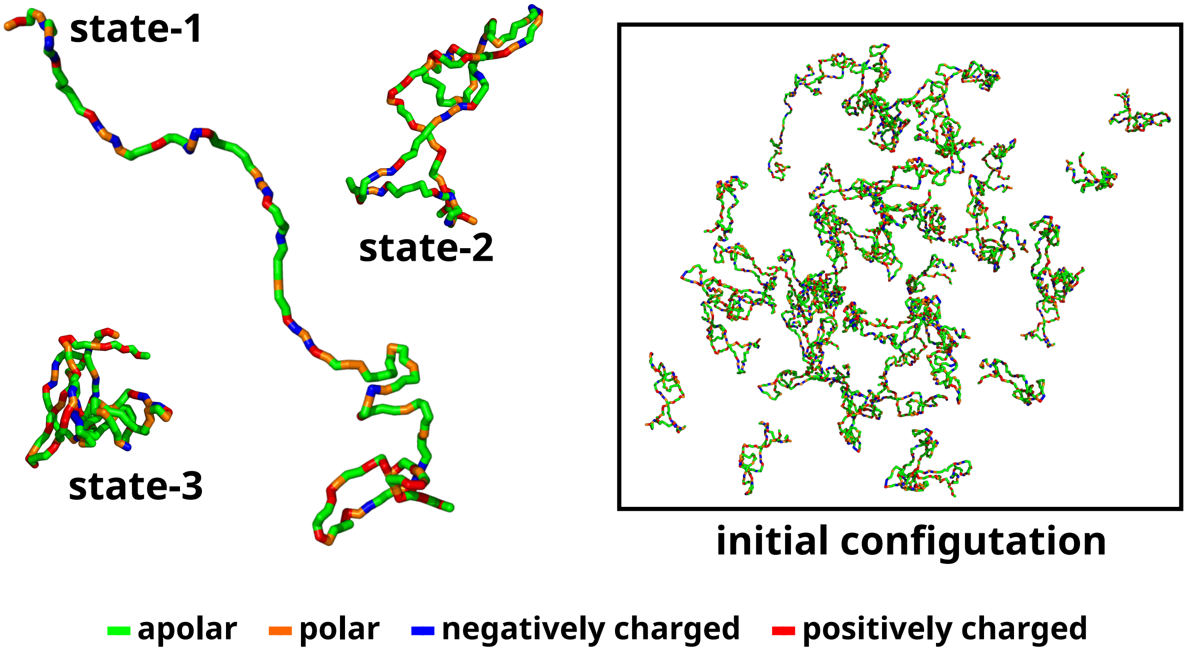


Figure S6: The left side of the figure shows the coarse-grained representation of three different conformation of $\alpha$S. state-1 is the most extended conformation, followed by state-2 and finally state-3 which is the most compact conformation. The right side of the figure shows the mixture of all these conformations with a total of 50 chains in a cubic box. The residues have been colour coded on the basis of their polarity/charge.

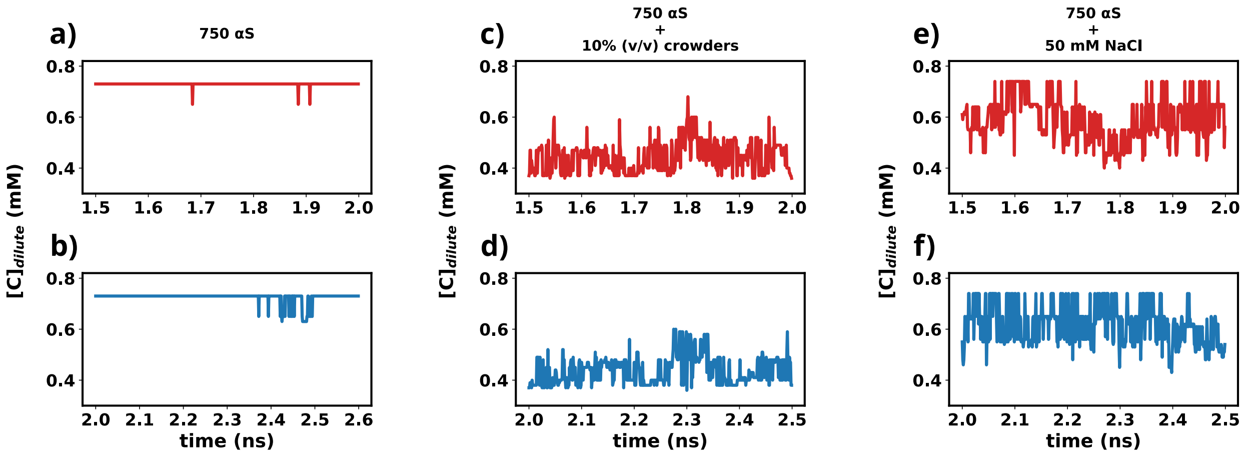


Figure S7: **a)** Time profile of the concentration of protein in the dilute phase between 1.5-2.0 $\mu$s. **b)** Time profile of the concentration of protein in the dilute phase between 2.0-1.5 $\mu$s. **c)** Time profile of the concentration of protein in the dilute phase between 1.5-2.0 $\mu$s in the presence of crowders. **d)** Time profile of the concentration of protein in the dilute phase between 2.0-1.5 $\mu$s in the presence of crowders. **e)** Time profile of the concentration of protein in the dilute phase between 1.5-2.0 $\mu$s in the presence of salt. **f)** Time profile of the concentration of protein in the dilute phase between 2.0-1.5 $\mu$s in the presence of salt.

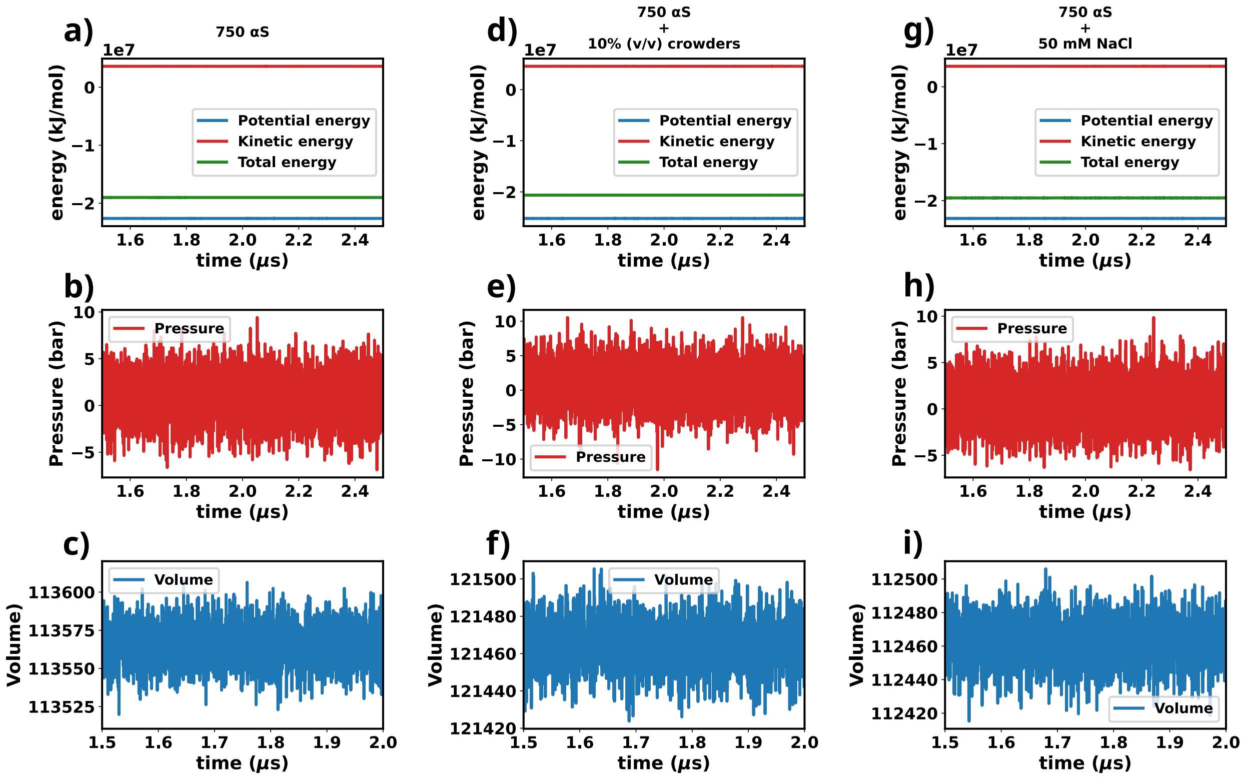


Figure S8: **a)** time profile of kinetic, potential and total enegies of the whole system for $\alpha$S in water. **b)** time profile of pressure of the whole system for $\alpha$S in water. **c)** time profile of simulation box volume for $\alpha$S in water. **d)** time profile of kinetic, potential and total enegies of the whole system for $\alpha$S in water in the presence of crowders. **e)** time profile of pressure of the whole system for $\alpha$S in water in the presence of crowders. **f)** time profile of simulation box volume for $\alpha$S in water in the presence of crowders. **g)** time profile of kinetic, potential and total enegies of the whole system for $\alpha$S in water in the presence of salt. **h)** time profile of pressure of the whole system for $\alpha$S in water in the presence of salt. **i)** time profile of simulation box volume for $\alpha$S in water in the presence of salt.

(3) Hung T Nguyen, Naoto Hori, and D Thirumalai. Condensates in rna repeat sequences are heterogeneously organized and exhibit reptation dynamics. *Nature Chemistry*, pages

1–11, 2022. 3
